# Drosophila ZDHHC8 palmitoylates scribble and Ras64B and controls growth and viability

**DOI:** 10.1101/365247

**Authors:** Katrin Strassburger, Evangeline Kang, Aurelio A. Teleman

**Affiliations:** German Cancer Research Center (DKFZ), 69120 Heidelberg, Germany; Heidelberg University, 69120 Heidelberg, Germany

**Keywords:** Drosophila, palmitoylation, CG34449, Ras64B, palmitoylome, ZDHHC8, scribble

## Abstract

**Summary:** Palmitoylation is an important posttranslational modification regulating diverse cellular functions. Consequently, aberrant palmitoylation can lead to diseases such as neuronal disorders or cancer. In humans there are roughly one hundred times more palmitoylated proteins than enzymes catalyzing palmitoylation (palmitoyltransferases). Therefore, it is an important challenge to establish the links between palmitoyltransferases and their targets. From publicly available data, we find that expression of human ZDHHC8 correlates significantly with cancer survival. To elucidate the organismal function of ZDHHC8, we study the Drosophila ortholog of hZDHHC8, CG34449/dZDHHC8. Knockdown of dZDHHC8 causes tissue overgrowth while dZDHHC8 mutants are larval lethal. We provide a list of 159 palmitoylated proteins in Drosophila and present data suggesting that scribble and Ras64B are targets of dZDHHC8.

## Introduction

Next generation sequencing technology has allowed the broad sequencing of tumors and adjacent healthy tissues to identify cancer-linked somatic mutations and cancer-linked genes. Many of these genes, however, are functionally uncharacterized and a role in cancer development has not been tested. This would be necessary to distinguish cancer ‘driver mutations’ from ‘passenger mutations’ which occur by chance and are neutral for the cancer phenotype. Drosophila is a good model organism for understanding gene function as the generation of loss-of -function mutants is fast, cancer-like phenotypes can be easily assayed, and flies have less genetic redundancy compared to humans, allowing the easier identification of phenotypes.

We focus here on a protein belonging to the family of palmitoyltransferases, which can be recognized in part due to their zinc finger DHHC domain (ZDHHC). ZDHHC enzymes catalyze the posttranslational, covalent attachment of the long chain acyl chains palmitate (C16:0) and stearate (C18:0) to cysteines of target proteins (S-palmitoylation). The binding of C16:0 to cysteines occurs via thioester linkages and can be reverted by thioesterases. Since S-palmitoylation is reversible and dynamic it can control a number of cellular processes such as subcellular trafficking, protein localization and stability as well as enzyme activity [1–3]. In mammals 23 proteins of the DHHC family have been identified as well as roughly 2000 palmitoylated proteins [4]. Therefore, it is an important challenge to understand which palmitoyltransferase targets which substrate.

Aberrant palmitoylation has been linked to many diseases ranging from neuronal disorders [5, 6] to cancer [4, 7–10]. Well known oncogenes whose activity is controlled by palmitoylation are for instance H-Ras and N-Ras proteins [11]. Recently, epidermal growth factor receptor (EGFR) has been shown to be palmitoylated by DHHC20 and loss of palmitoylation leads to increased EGFR signaling, cell migration and transformation [12]. DHHC3 was shown to have growth promoting effects in breast cancer cells regulating oxidative stress and senescence [13]. Consequently, palmitoyltransferases are being pursued as targets for therapeutic intervention in cancer [14].

We show here that knockdown of the Drosophila orthologue of human ZDHHC8 (CG34449 or dZDHHC8) causes tissue overgrowth. dZDHHC8 mutants are larval lethal and have metabolic phenotypes. We identify potential targets of dZDHHC8 and suggest that dZDHHC8 loss of function phenotypes partly arise from decreased palmitoylation of scribble and Ras64B proteins.

## Results

### Drosophila ZDHHC8 knockdown causes tissue overgrowth

We aimed to functionally characterize novel cancer relevant genes. To identify such genes we searched through publicly available databases for genes whose expression correlates with patient survival. In The Protein Atlas database (www.proteinatlas.org) [15] we found that the expression of ZDHHC8 correlates with cancer prognosis (Supplemental Figure 1A). In renal and cervical cancer, high expression of ZDHHC8 is significantly correlated with shorter survival times, whereas in lung and pancreatic cancers high expression significantly correlates with longer survival rates (Supplemental Figure 1A). These data suggest that ZDHHC8 may play a role in cancer incidence and/or progression, but that this role likely depends on tumor type or context. Since ZDHHC8 has not been previously linked to these cancers we decided to study ZDHHC8 function, and whether it affects tissue growth or cell proliferation.

ZDHHC8 is a member of the palmitoyl-transferase group that is highly conserved across evolution. BLASTing the protein sequence of human ZDHHC8 against the Drosophila proteome identifies the uncharacterized gene CG34449 as a top hit with an E-value of 10^−76^ (Supplemental Figure 1B). Conversely, a BLAST search of the human proteome using CG34449 protein sequences identifies ZDHHC8 as the top hit with an E-value of 10^−87^ (Supplemental Figure 1B). Hence, CG34449 is the Drosophila orthologue of hZDHHC8. We therefore used Drosophila melanogaster as a model system to study CG34449 function *in vivo* and refer to it as dZDHHC8.

We first tested whether dZDHHC8 regulates tissue growth in the Drosophila wing, since the wing is flat and its size is easily quantified. The wing consists of two cellular layers, a dorsal and a ventral layer (Figure 1A). Unbalanced growth of the two layers leads to bending of the wing. We knocked down dZDHHC8 with three independent RNAi lines, targeting different regions of the gene in the dorsal part of the wing using the apterous-Gal4 driver (apG4) (Figure 1A). In all three cases we observed a downward bending of the wings, indicating that knockdown of dZDHHC8 leads to tissue overgrowth (Figure 1A). Since we observe this phenotype with three independent dsRNA’s, this makes it highly unlikely that off-target effects are responsible for the phenotype. Also when dZDHHC8 is knocked down in the posterior part of the wing using the engrailed-Gal4 driver, (enG4>dZDHHC8 RNAi #1, Figure 1B) the ratio of posterior to anterior wing size increases mildly (7%) but significantly compared to control wings (enG4>+, Figure 1B). To find out whether this increased tissue size is due to more cells or larger cells, we quantified cell size in the posterior compartment where dZDHHC8 was knocked down and normalized it to cell size in the control anterior compartment. This was done by counting the number of cells via trichomes in a region of defined size, and then calculating the ratio of area per cell. We found that cell size was not affected by dZDHHC8 knockdown (enG4>dZDHHC8 RNAi, Figure 1C) when compared to control wings (enG4>+, Figure 1C). This suggests that the increased tissue size is due enhanced cell proliferation upon dZDHHC8 knockdown. Knocking down dZDHHC8 ubiquitously using tubulin-Gal4 (tubG4>GFP, dZDHHC8 RNAi, Figure 1D) often resulted in extra vein material, which could not be observed in control wings (tubG4>GFP, Figure 1D).

**Figure 1:**
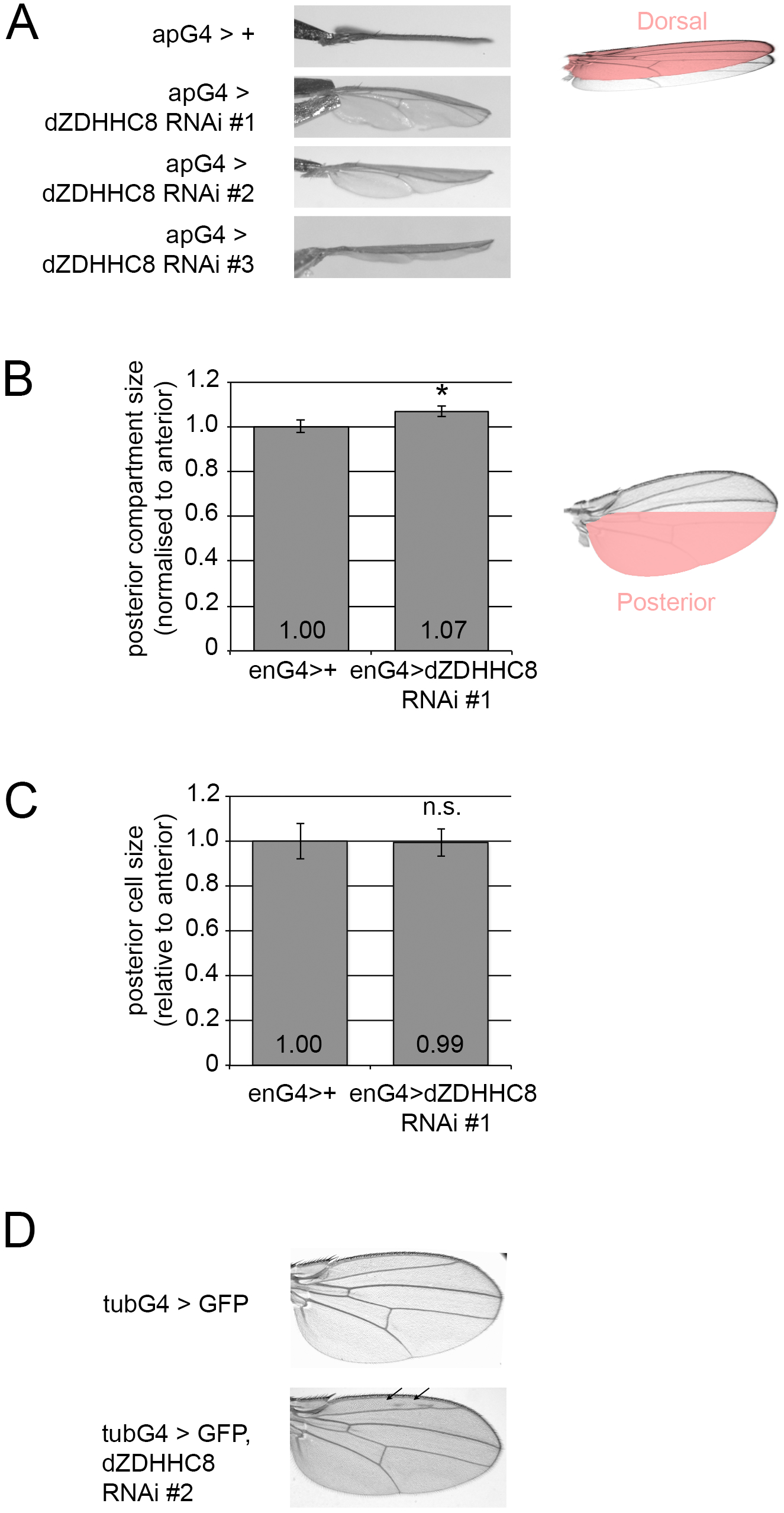
dZDHHC8 knockdown causes tissue overgrowth due to increased cell proliferation. **(A)** Three independent RNAi lines targeting different regions of dZDHHC8 mRNA result in tissue overgrowth and downward bending of the wing when expressed in the dorsal compartment using apterous-Gal4 (apG4) at 25°C. **(B)** Expression of dZDHHC8 RNAi in the posterior compartment of the wing using engrailed-Gal4 (enG4>dZDHHC8 RNAi #1) increases posterior compartment size normalized to anterior when compared to control wings (enG4>+). Error bars = stdev. n≥9. ^∗^ ttest=2×10^−5^ **(C)** The size of wing cells is not altered upon dZDHHC8 knockdown suggesting that tissue overgrowth is caused by enhanced cell proliferation. dZDHHC8 is knocked down in the posterior compartment with engrailed-Gal4 (enG4>dZDHHC8 RNAi #1). Cell size was determined by counting the number of cells (via trichomes) within a wing area of defined size. Error bars = stdev. n≥9. **(D)** Ubiquitous expression of dZDHHC8 RNAi using tubulin-Gal4 (tubG4>GFP, dZDHHC8 RNAi #2) often results in formation of extra vein material.

### dZDHHC8 knockouts are larval lethal with metabolic phenotypes

To further study the function of dZDHHC8 we generated two different knockout alleles. Knockout line 1 (KO1) lacks most of the dZDHHC8 genomic sequence, including CG34450, which is annotated as a separate gene within dZDHHC8 in Flybase (Figure 2A) [16]. Since these two genes were previously annotated in Flybase as one linked gene and split into two genes in release 5.2 of the genome annotation, we tested whether they are indeed independent of each other. We knocked down CG34450 in Drosophila S2 cells and measured levels of dZDHHC8 mRNA by qRT-PCR using oligos that anneal to different regions of dZDHHC8 (Supplemental Figure 2). Knockdown of CG34450 caused mRNA levels of both CG34450 and dZDHHC8 to drop (Supplemental Figure 2). Transcript levels of dZDHHC8 decreased less than levels of CG34450 transcript, although this could be explained by the fact that dZDHHC8 has multiple alternatively-spliced transcript isoforms (Figure 2A). This indicates that mRNA levels of dZDHHC8 and CG34450 are linked in some way and perhaps they are not separate genes. In a second knockout line we removed a small genomic region common to all isoforms of dZDHHC8 including the catalytic domain (KO2 Figure 2A). We confirmed that dZDHHC8 knockouts do not have dZDHHC8 protein by western blot analysis using a dZDHHC8 antibody which we raised in guinea pigs (Figure 2B). (As is often the case for membrane-integral proteins, ZDHHC8 forms multiple bands on an SDS-PAGE gel.)

**Figure 2:**
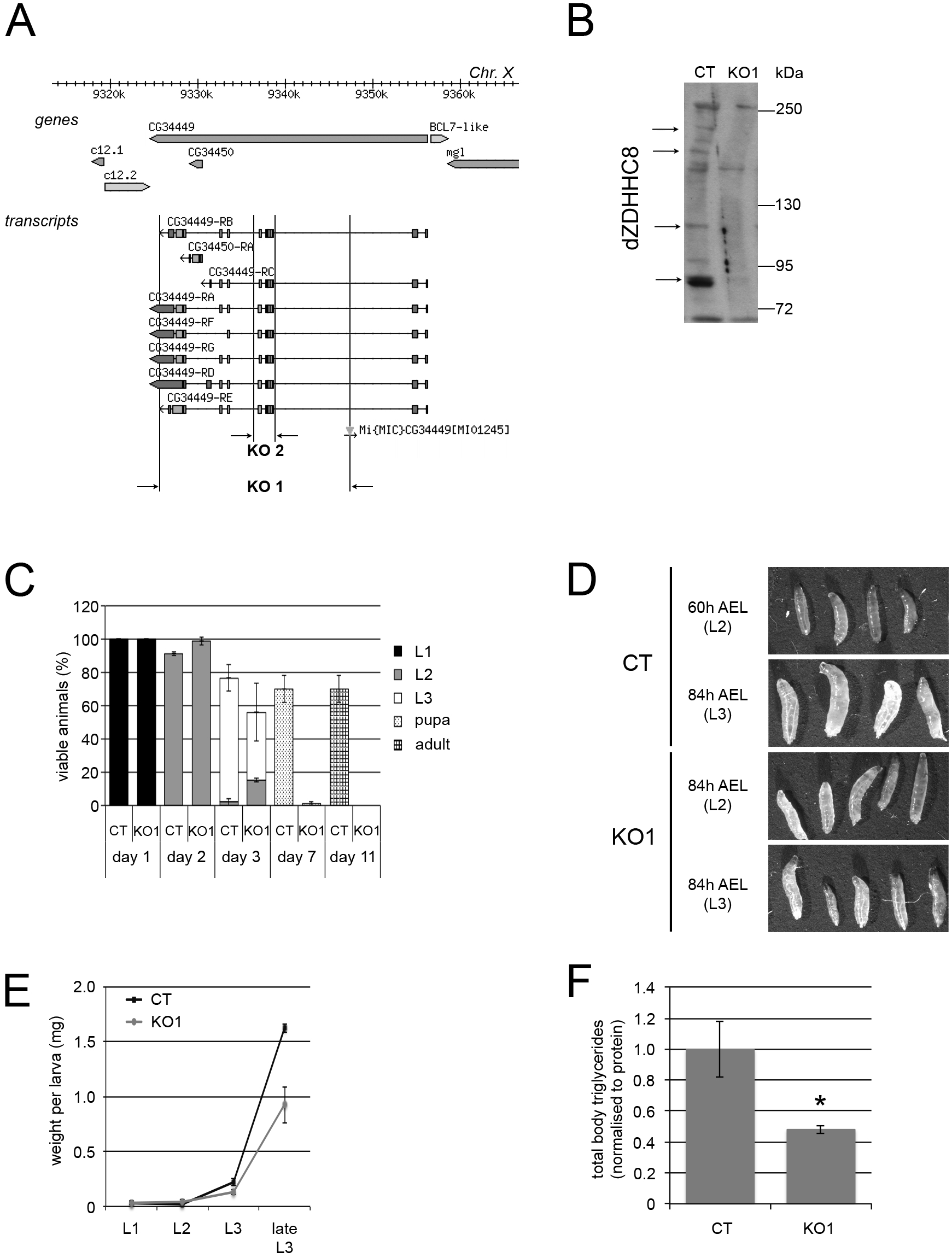
dZDHHC8 knockout phenotypes. **(A)** Schematic of the dZDHHC8 genomic locus, indicating transcript isoforms and knockout regions. Two independent dZDHHC8 knockout lines were generated: knockout 1 (KO1) which removes the majority of dZDHHC8 as well as CG34450, and knockout 2 (KO2) which removes a small region common to all dZDHHC8 isoforms including the catalytic domain. **(B)** dZDHHC8 knockout larvae do not have dZDHHC8 protein. dZDHHC8 protein levels were analysed by immunoblotting extracts from control (CT) or dZDHHC8 knockout (KO1) larvae, separated on a 6% gel with a dZDHHC8 specific antibody. dZDHHC8 appears as multiple high molecular weight bands. Arrows indicate dZDHHC8 protein absent in KO1. **(C)** dZDHHC8 mutants are larval lethal. Control (CT) and dZDHHC8 knockouts (KO1) were collected as L1 larvae and the number and developmental stage (judged by mouth hook morphology) of viable animals was determined after 1-11 days. KO1 animals start dying at the L2/L3 transition with very few animals reaching pupal stage. Some lethality of control animals is observed in this experiment, due to all animals being picked out of the food and sorted daily for scoring. Error bars = stdev. n=3×60 L1/genotype. **(D)** dZDHHC8 phenotypes appear at the L2/L3 transition. Shown are control (CT) and dZDHHC8 mutant (KO1) larvae imaged at 60h after egg laying (AEL) (corresponds to wildtype larval stage 2) or 84h AEL (corresponds to wildtype larval stage 3). The developmental stage of the depicted larvae, judge by the mouth hook morphology is indicated. KO1 larvae are limited in their capability to molt from L2 to L3, either growing bigger for a few days as L2, or molting but never reaching the size of control L3 larvae. **(E)** dZDHHC8 mutants weigh less than controls. dZDHHC8 knockout larvae (KO1) are lighter than control (CT) larvae with the difference starting at the L2/L3 transition. Error bars = stdev. n=3×20 larvae/genotype. **(F)** dZDHHC8 mutants are lean. Shown are total body triglyceride levels, normalized to total body protein, for control (CT) and dZDHHC8 knockout (KO1) L3 larvae. Error bars = stdev. n=3×6 larvae/genotype. ^∗^ ttest=0.04.

Both dZDHHC8^KO1^ and dZDHHC8^KO2^ animals do not reach adulthood, but die as larvae. To determine precisely at which stage dZDHHC8^KO1^ animals die, we collected control (CT) or dZDHHC8^KO1^ newly-hatched first-instar larvae and examined the number and developmental stage of live animals during the following 11 days (Figure 2C). In this and all other experiments, control animals are w^1118^ animals with the same genetic background as the dZDHHC8^KO1^ (see Materials & Methods). This revealed that dZDHHC8^KO1^ animals die around the transition from larval stage 2 to 3. This is also true for dZDHHC8^KO2^ animals (not shown). Due to the similarity in phenotype of the two alleles, we subsequently characterized in depth allele KO1. Examination of animal morphology showed that dZDHHC8 knockouts have a defect in molting from L2 to L3. Many animals fail to molt, and remain L2 for a few days, continuing to grow to a size larger than normal L2 larvae (Figure 2D). Other dZDHHC8^KO1^ larvae molt to L3, but fail to grow and do not reach the size of control animals (Figure 2D, bottom row). Occasionally, we observed dead L3 larvae with their L2 carcass still attached to the mouth hooks (not shown). dZDHHC8 knockout animals also have metabolic phenotypes that become apparent at the L2/L3 developmental stage. dZDHHC8^KO1^ weigh less and have less total body triglycerides than control animals (Figure 2E-F). Unexpectedly, we did not observe tissue overgrowth in dZDHHC8^KO1^ animals, but this may be due to differences in tissue-specific versus systemic loss of function, as also observed for other growth regulators (see Discussion). In sum, the data from knockdown and knockout experiments suggest a function of dZDHHC8 in growth and metabolism.

### dZDHHC8 resides at the Golgi

Palmitoyltransferases such as dZDHHC8 typically localize either to the ER or to the Golgi [17]. To determine the subcellular localization of dZDHHC8 we first tested whether our self-made dZDHHC8 antibody specifically detects endogenous dZDHHC8 by immunostaining. To this end, we stained Drosophila S2 cells that had been treated either with control dsRNA (luciferase dsRNA) or a dsRNA targeting dZDHHC8 (Figure 3A). This revealed a speckled staining in control cells which was absent in dZDHHC8 knockdown cells, suggesting that the antibody is specific for dZDHHC8. To figure out whether the speckled appearance is due to dZDHHC8 localization at a specific organelle, we expressed markers for different subcellular compartments in Drosophila S2 cells and co-stained for endogenous dZDHHC8 (Figure 3B). We found that dZDHHC8 co-localizes with a Golgi-tethered protein (Golgi-tethered Fringe-myc [18]).

**Figure 3:**
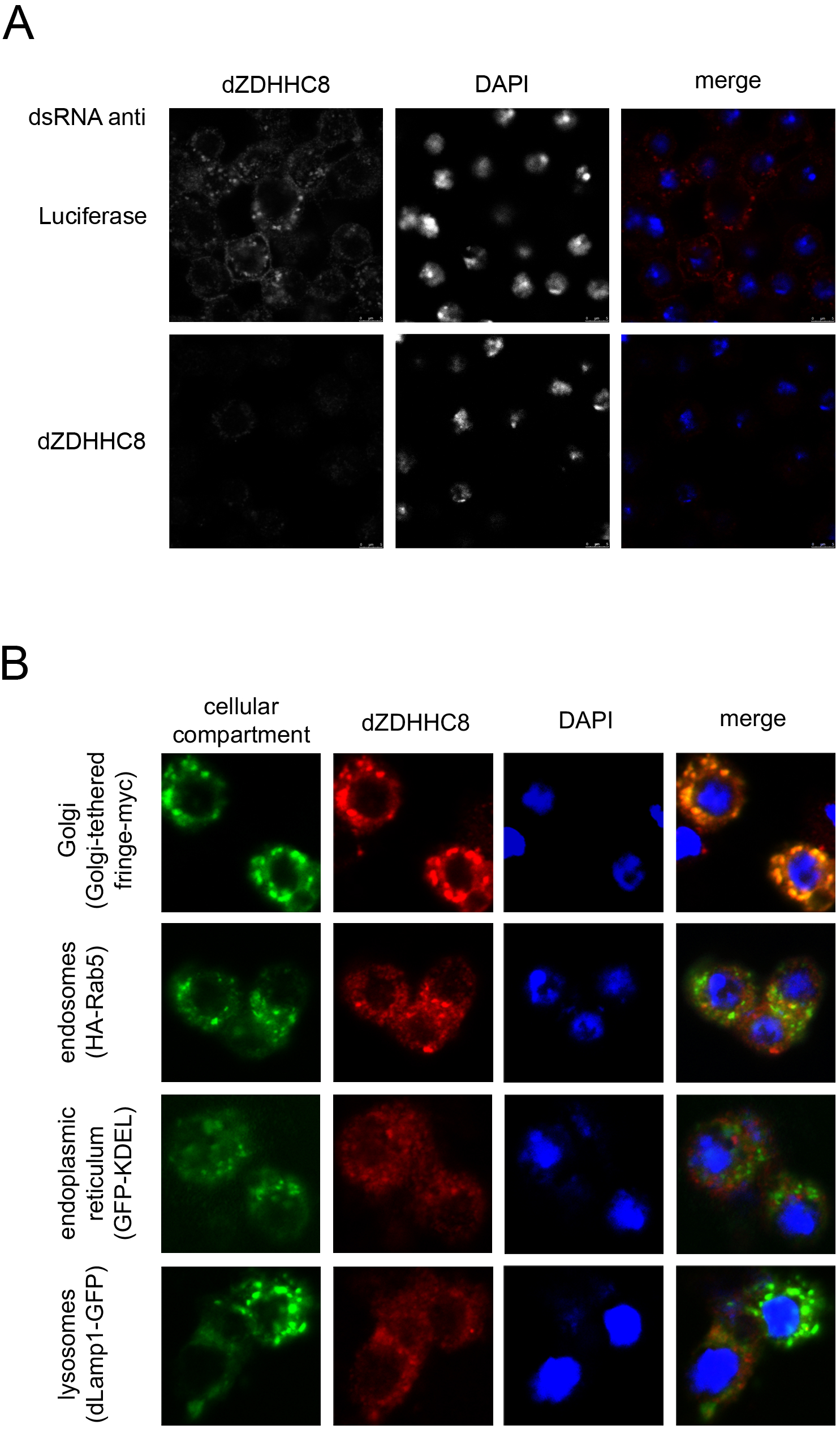
dZDHHC8 resides at the Golgi. **(A)** Antibody against dZDHHC8 specifically detects dZDHHC8 in immunofluorescent stainings of Drosophila S2 cells treated with control (luciferase) dsRNA as there is no staining upon dZDHHC8 knockdown (dsRNA anti dZDHHC8). **(B)** Endogenous dZDHHC8 (red) was co-stained with markers for different cellular compartments (green). Overexpressed organelle markers are Golgi-tethered fringe-myc, HA-Rab5 marking endosomes, GFP-KDEL marking endoplasmic reticulum and dLamp1-GFP marking lysosomes. DAPI was used to stain nuclei. dZDHHC8 co-localises with the Golgi-tethered marker.

### Acyl-Biotin exchange (ABE) assay identifies palmitoylated proteins in Drosophila as well as dZDHHC8 targets

To identify proteins that are palmitoylated by dZDHHC8 we used a method called acyl-biotin exchange (ABE). This method allows acyl chains on proteins to be replaced with biotin for streptavidin pull down (Figure 4A). Subsequently, acylated proteins can be identified by mass spectrometry (ABE-MS) or western blotting (ABE-WB, Figure 4A). As a negative control, the hydroxylamine (H_3_NO) is omitted, and acylated proteins are the ones observed differentially represented in the +H_3_NO pulldown compared to the -H_3_NO pulldown. We performed ABE-MS from lysates of control and dZDHHC8 knockout animals. Three independent experiments yielded a list of 159 genes that are significantly acylated in either the control or knockout animals or both (Supplemental Table 1, FDR<0.1). This provides a list of proteins that are acylated (e.g. palmitoylated, stearoylated, or myristoylated) in Drosophila and may constitute a useful resource for the community. Amongst these proteins are proteins previously known to be palmitoylated in mammals such as Flotilin 1 (Flo1) and Flotilin 2 (Flo2) or cysteine string protein (Csp). We further validated our mass spectrometry data by performing ABE-WB on lysates of larvae overexpressing HA-tagged Csp and Ras64B and confirmed that Csp and Ras64B are palmitoylated in Drosophila (Figure 4B).

**Figure 4:**
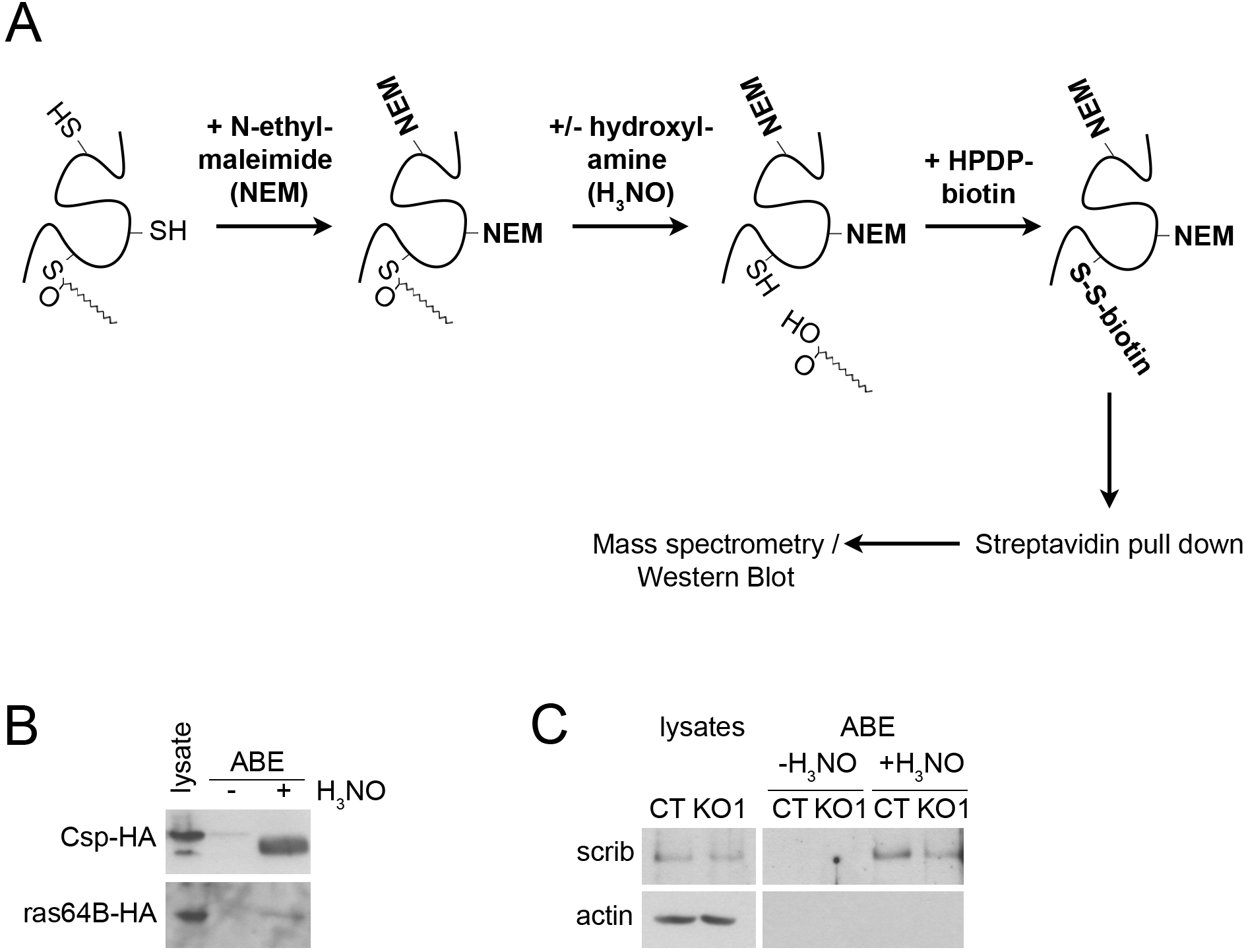
Acyl-Biotin Exchange (ABE) assay identifies acylated proteins in Drosophila as well as potential dZDHHC8 targets. **(A)** Schematic representation of an Acyl-Biotin Exchange (ABE) assay followed by mass spectrometry (ABE-MS) to identify acylated proteins (ie palmitoylated, stearoylated, etc). First, lysates are treated with N-ethylmaleimide to block free cysteine-residues on proteins. Acylation on proteins is then removed by hydroxylamine (H_3_NO) treatment yielding free cysteines that were previously acylated. Free cysteines subsequently react with thiol-reactive HPDP-biotin and biotinylated proteins are pulled down with strepavidin-beads. Eluted proteins are analysed by mass spectrometry or western blot (ABE-WB). Every ABE-assay includes a non-H_3_NO treated control to distinguish acylated proteins from unspecifically biotinylated proteins. **(B)** Csp and Ras64B are acylated. ABE assay was performed on lysates from L3 larvae expressing HA-tagged Csp or Ras64B under the control of heatshock-Gal4 24h after heatshock. Csp and Ras64B are acylated as they are specifically pulled down in the H_3_NO treated, but not in H_3_NO untreated control samples. **(C)** Scrib is palmitoylated, partly by dZDHHC8. Shown are endogenous scrib protein levels from control (CT) and dZDHHC8 knockout (KO1) larvae in lysates and after ABE assay. Scrib is detected after ABE in H_3_NO treated samples of CT and to a lesser extend of KO1 larvae, but not in control samples (-H_3_NO).

To identify dZDHHC8 targets we screened our ABE-MS data for proteins that are less palmitoylated in the dZDHHC8^KO1^ samples compared to the control samples (Supplemental Table 2). With a cutoff of 40% decrease in palmitoylation in KO compared to control, we identified 13 proteins as potential targets of dZDHHC8, however the difference in control vs KO was not statistically significant.

To link the dZDHHC8 loss of function fly phenotype to a dZDHHC8 target protein we initially focused on scribble because of its known tumor suppressor function. First, we performed ABE-WB of endogenous scribble from control and knockout larvae. We found that scribble is palmitoylated at least in part by dZDHHC8 as there is less palmitoylated scribble pulled down in knockout animals compared to control animals (Figure 4C, compare KO1 +H_3_NO to CT +H_3_NO). This suggests that even though the mass spectrometry analysis did not yield statistically significant differences between control and dZDHHC8^KO1^ samples, the data can still be used as a list of potential dZDHHC8 targets.

We hypothesized that impaired palmitoylation of scribble could lead to changes in scribble activity, which we tested in three different ways: (1) To test if dZDHHC8 lethality was due to impaired scribble activity we tested whether removing one copy of scrib in a dZDHHC8^KO1/+^ heterozygous mutant background would lead to synthetic lethality. However, dZDHHC8^KO1/+^, scrib^-/+^ transheterozygous mutants were as viable as their control counterparts (data not shown). (2) Scribble regulates cell polarity in a protein complex with lethal giant larvae (Lgl) and discs large (Dlg), and follicle cells lacking scribble are rounded and often multilayered [19]. To examine whether dZDHHC8 loss-of-function leads to defects in follicle cell polarity we stained dZDHHC8 mutant follicle clones for Dlg (Supplemental Figure 3A). However, follicle cells lacking dZDHHC8 appeared morphologically normal as a monolayer and we could not observe any differences in Dlg subcellular localization in dZDHHC8 knockout clones compared to neighboring cells (Supplemental Figure 3A). (3) Scribble also affects Hippo signaling [20]. We tested whether the activity of Yorkie, a transcriptional co-activator downstream of Hippo was affected using the transcriptional reporter expanded-lacZ (ex-lacZ). We generated dZDHHC8 knockout clones in the wing disc carrying ex-lacZ and immuno-stained them with an antibody against β-galactosidase (Supplemental Figure 3B). Again, we did not see any differences in expanded expression in knockout clones compared to neighboring wildtype cells (Supplemental Figure 3B). Even though dZDHHC8 seems to palmitoylate scribble, we could not find evidence that the degree of drop in scribble palmitoylation caused by dZDHHC8 loss is sufficient to affect scribble activity. Therefore, we focused on a different target of dZDHHC8.

### dZDHHC8 affects Ras64B stability

Looking at the list of potential dZDHHC8 targets our attention was caught by Ras64B because ras proteins are known to play a role in growth control. We looked for genetic interaction of dZDHHC8 and Ras64B and found that overexpression of Ras64B enhances dZDHHC8 mutant lethality (Figure 5A). To verify our ABE-MS data we performed an ABE-WB assay from larval lysates of Ras64B-HA overexpressing control and dZDHHC8 knockout animals (Figure 5B). We found that knockout animals have indeed less palmitoylated Ras64B (Figure 5B, ABE KO1 + H_3_NO versus ABE CT + H_3_NO) however total Ras64B protein was also decreased to a similar extent (Figure 5B, first two lanes). This suggests that loss of dZDHHC8 either leads to reduced Ras64B palmitoylation and, as a consequence, reduced Ras64B stability, or the other way around - that dZDHHC8 affects Ras64B stability, and as a consequence palmitoylation. We therefore studied the effects of dZDHHC8 on Ras64B stability and palmitoylation. We tested whether dZDHHC8 loss of function leads to decreased Ras64B stability by expressing Ras64B-HA in control and dZDHHC8^KO1^ animals (Figure 5C). We used a heat-inducible driver (hs>Ras64B) and looked at Ras64B-HA protein levels at different time points after heat-shock. This showed that Ras64B-HA stability is strongly dependent on dZDHHC8 with Ras64B-HA protein levels being decreased in dZDHHC8 knockout animals even when Ras64B-HA is expressed at low levels without any heat-shock (Figure 5C, 0h post hs). Ras64B has five cysteines that could serve as potential palmitoylation sites. Two cysteines belong to the c-terminal CAAX motif (CCLM amino acids 189-192), which is required for ras farnesylation, which in turn serves as a priming event for ras palmitoylation [21]. As expected, mutation of cysteines in the CAAX motif resulted in a complete loss of Ras64B palmitoylation (Figure 5G). To test whether any of the other three cysteines is palmitoylated we mutated each of them to alanine and performed ABE-WB on lysates of larvae expressing Ras64B wildtype (WT) or Ras64B alanine mutants (C46A, C120A, C147A, Figure 5D). To our surprise, each of the alanine mutant versions is less palmitoylated compared to WT Ras64B (compare +H_3_NO lanes, Figure 5D) yet each mutant is still palmitoylated (compare + to - H_3_NO for each mutant). This suggests that Ras64B is palmitoylated on more than one site. Therefore, we mutated two cysteines at the same time (Figure 5E) and found that again palmitoylation is present in each of the three double-mutants (Figure 5E), suggesting that all three sites are palmitoylated. In agreement with this, simultaneous mutation of all three cysteines to alanine completely abolished palmitoylation of Ras64B (Figure 5F). In sum, Ras64B is palmitoylated on Cys46, Cys120 and Cys147.

**Figure 5:**
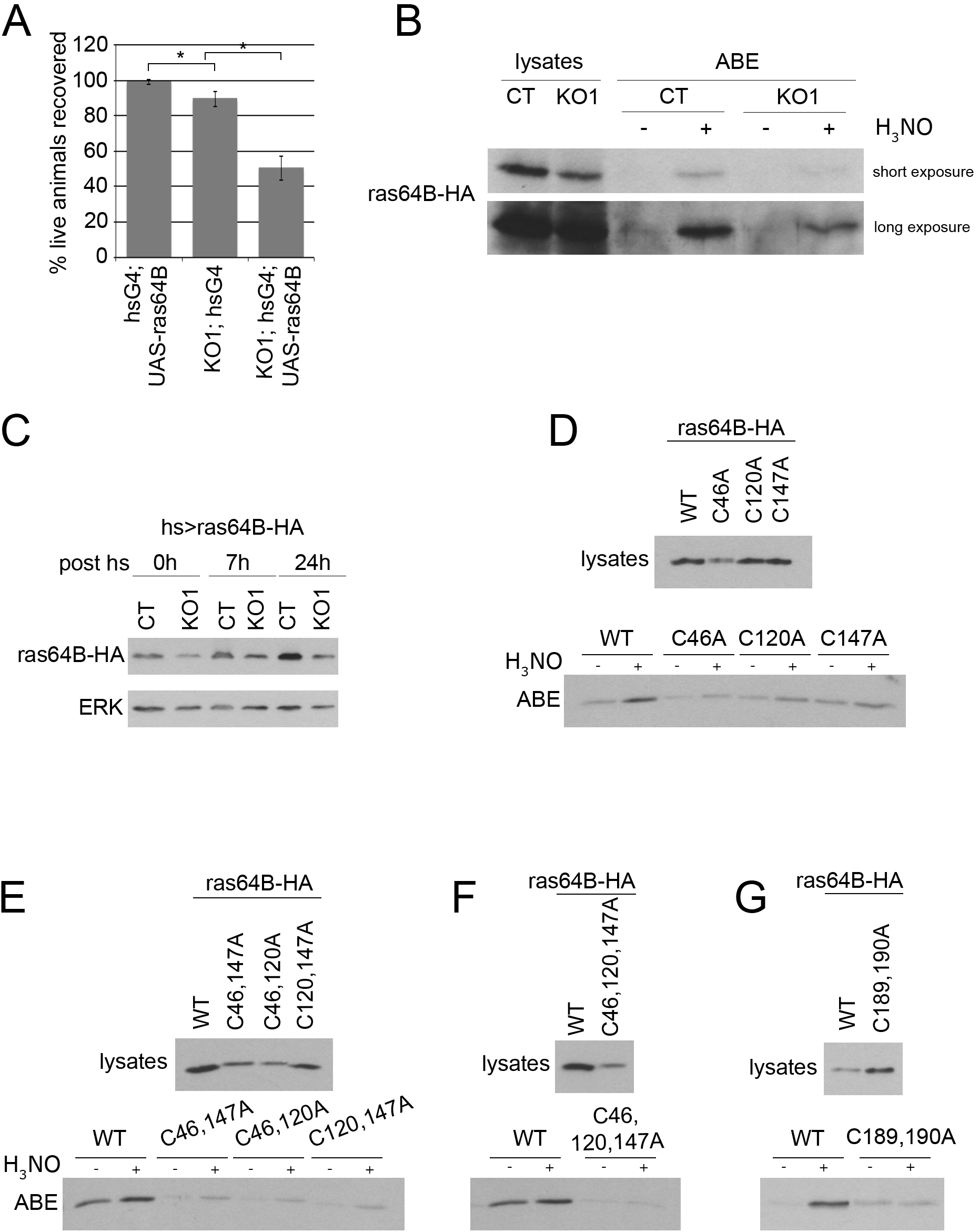
dZDHHC8 affects Ras64B stability. **(A)** Ras64B overexpression decreases viability of dZDHHC8 knockouts. Four days after L1 collection, the number of live Ras64B overexpressing larvae (hsG4; UAS-Ras64B), dZDHHC8 homozygous mutant larvae (KO1; hsG4) or larvae overexpressing Ras64B in dZDHHC8^KO1^ mutant background (KO1; hsG4; UAS-Ras64B) was determined. Ras64B overexpression in the dZDHHC8^KO^ mutant background causes synergistic lethality. Error bars = stdev. n=3×60 larvae/genotype. ^∗^ ttest≤0.04. **(B)** ABE assay was performed on lysates from control (CT) and dZDHHC8 knockout (KO1) early L3 larvae expressing Ras64B under the control of heatshock-Gal4 24h after heatshock. KO1 larvae have less palmitoylated Ras64B-HA when compared to WT controls but also less total Ras64B-HA protein. **(C)** dZDHHC8 affects Ras64B stability. Protein levels of Ras64B-HA expressed with hsG4 in control (CT) and dZDHHC8 knockout (KO1) larvae were assayed by immunoblotting at different time points after heat-shock. ERK was used as a loading control. KO1 larvae have less Ras64B-HA protein than CT larvae at any time point. **(D)** Ras64B is palmitoylated on at least two cysteines. Larvae expressing HA-tagged wildtype Ras64B, or mutant Ras64B where each of the three cysteine residues present in Ras64B is mutated to alanine. All expression constructs are inserted in the same genomic locus using phiC31-directed integration, and their expression induced using hsG4 and collecting samples for ABE 24h after heat-shock. All Ras64B-HA C/A mutants are still palmitoylated, albeit less than WT Ras64B-HA. Note that Ras64B C46A protein levels are lower than wildtype Ras64B. **(E-G)** Ras64B is palmitoylated on all three cysteines C46, C120, C147. Assay performed as in (B). Each of the Ras64B-HA double cysteine mutants is still palmitoylated (E). Only mutating all three cysteines at the same time (F) or mutating the cysteine before and within the CAAX motif (G) abolishes palmitoylation. Note that protein levels of Ras64B versions carrying C46A are lower.

Studying the various Ras64B mutants we realized that whenever Cys46 is mutated to alanine (Figure 5D-F, C46A) Ras64B protein levels strongly drop in the lysates, just like they do when WT Ras64B is expressed in dZDHHC8 mutants (Figure 5B). This suggests that Cys46 plays a role in Ras64B protein stability. Until now, however, it is unclear whether dZDHHC8 palmitoylates Ras64B at Cys46 thereby regulating its stability, or whether dZDHHC8 affects Ras64B stability indirectly through some other mechanism.

## Discussion

The function of hZDHHC8 has been mainly characterized to date in the brain, where loss of the genomic hZDHHC8 locus in microdeletions at 22q11 is associated with cognitive deficits and schizophrenia [22–24]. To our knowledge, hZDHHC8 has not been implicated in cancer so far, except that knockdown of hZDHHC8 makes mesothelioma cells more sensitive to radiotherapy [25]. We report here the functional characterization of the Drosophila ortholog of hZDHHC8, CG34449/dZDHHC8. Knockdown of dZDHHC8 in the Drosophila wing resulted in tissue overgrowth due to increased cell proliferation (Figure 1) suggesting that dZDHHC8 inhibits cell proliferation tissue autonomously. When we generated knockout flies, we found that dZDHHC8 mutant flies are small and lean and die as larvae (Figure 2). The lethality may be the result of loss of palmitoylation on all dZDHHC8 targets and is therefore very pleiotropic. The fact that mild knockdown results in tissue overgrowth whereas complete loss of function is lethal has been reported also for other genes, as for instance TSC2 loss of function clones in the eye cause overgrowth whereas ubiquitous loss of function of TSC2 is embryonic lethal [26].

In their characterization of palmitoyltransferases and thioesterases, Bannan and colleagues previously showed that most of Drosophila palmitoyltransferases localize to the ER or Golgi [17]. Our findings complement their findings by showing that also dZDHHC8 localizes to the Golgi (Figure 3). Furthermore, we identify here palmitoylated proteins on a proteome wide level in Drosophila by performing ABE assays from larvae. This yields a list of 159 acylated target proteins (Supplemental Table 1) which will hopefully constitute a useful resource for the community.

Comparing palmitoylated proteins of control and dZDHHC8 mutant larvae we find scribble to be palmitoylated in part by dZDHHC8. Scribble palmitoylation is decreased in dZDHHC8 mutants, however not completely abolished (Figure 4). This suggests that there is another palmitoyltransferase acting on scribble. Potentially, this could be the fly homologue of ZDHHC7, as human ZDHHC7 has been shown to palmitoylate scribble in HEK293A cells [27]. In addition to scribble, we also verified Ras64B as a target of dZDHHC8 (Figure 5). We find that dZDHHC8 lethality is enhanced by overexpression of Ras64B suggesting that they act synergistically (Figure 5A). The mechanism appears rather complicated as loss of dZDHHC8 leads to destabilization of Ras64B. It has been proposed for other proteins that lack of palmitoylation can lead to their destabilization [28, 29]. In this case here it is unclear whether Ras64B destabilization is due to impaired palmitoylation or whether dZDHHC8 acts on Ras64B in a different manner to affect its stability, and as a consequence we also see reduced palmitoylation. It has been reported that overexpression of Ras64B in the wing causes the formation extra vein material [30]. Interestingly, we find that knockdown of dZDHHC8 also leads to ectopic veins (Figure 1D) again pointing towards dZDHHC8 acting on Ras64B.

We find that knockdown of dZDHHC8 causes tissue overgrowth, which is a cancer-relevant phenotype. Furthermore, we provide data suggesting that scribble and Ras64B are targets of Drosophila ZDHHC8. These findings possibly shed some light on the epidemiological links between hZDHHC8 and cancer (Supplemental Figure 1). Admittedly, the relationship between hZDHHC8 and cancer progression is complex, because in some cases elevated ZDHHC8 expression correlates with poor prognosis, and in some cases it correlates with good prognosis (Supplemental Figure 1). Hence significant more work will be required to understand this relationship. Nonetheless, the fact that dZDHHC8 knockdown causes a cancer-relevant phenotype in Drosophila suggests it may be worthwhile looking at hZDHHC8 more closely.

## Materials & Methods

### Plasmids and fly strains

The plasmids used in the study are listed in Supplemental Table 3. Full sequences can be obtained upon request. Sequences of oligos are in Supplemental Table 4. apterous-Gal4, engrailed-Gal4, tubulin-Gal4, heatshock-Gal4, UAS-GFP, Mi{MIC}dZDHHC8^MI01245^, UAS-ras64B RNAi (stock 29318), and hsFlp, hs-SceI/Cyo, y,w, nos-phiC31(X); Sco/Cyo (stock 34770) were obtained from Bloomington Stock Center. UAS-dZDHHC8 RNAi lines GD5521 (#1) and GD48025 (#2) were obtained from Vienna Drosophila Research Center. From DGRC we obtained expression constructs for Csp-HA (clone UFO08205) and ras64B-HA (clone UFO6356). For expression of tagged ras64B in Drosophila, plasmids pKH410, 411, 412, 413, 414, 430, 433, 434 and 437 (Supplemental Table 3) were injected into vk33 (Bloomington stock 32543) flies to allow site directed insertion on the third chromosome. The dZDHHC8 RNAi #3 line was generated by injecting pKH364 into w^1118^ flies.

### dZDHHC8 knockout generation

dZDHHC8 knockout 1 was generated using the MIMIC line Mi{MIC}dZDHHC8^MI01245^, in which the MIMIC cassette flanked by attP sites was replaced by Recombination Mediated Cassette Exchange (RMCE) with a knockout cassette flanked by attB sites [31]. The knockout cassette consisted of a yellow marker gene, followed by a SceI restriction site, a sequence homologues to the genomic region downstream of the knockout locus and a mini white gene (pKH304). RMCE was achieved by injecting pKH304 (Supplemental Table 3) into F1 embryos of nos-phiC31 × Mi{MIC}dZDHHC8^MI01245^ and screening for w+, y+ knockout donors. The correct orientation of the knockout cassette in the donor line was verified by PCR. The donor line was subsequently crossed to hs-SceI, and SceI expression was induced by heat-shocking the F1 on day 2 for 1h at 37°C. 600 F1 virgins that were w+ and mosaic for yellow were crossed to balancer males and the F1 was screened for w+, y-progeny. In those animals the double-strand break induced by SceI restriction had been repaired by homologous recombination using the genomic sequence of the knockout cassette. This led to loss of the yellow marker gene. We recovered 1 female from all progeny of the 600 crosses. dZDHHC8 knockout 2 was generated by homologous recombination as described previously [32] using pKH261 as a donor vector.

### Back-crossing of dZDHHC8 mutants

To clean the genetic background of dZDHHC8 mutants, heterozygous virgin females carrying the dZDHHC8^KO^ allele were back-crossed to w^1118^ males for 4 generations. In parallel FM7 balancer virgins were back-crossed to w^1118^ males to place the FM7 balancer into the w^1118^ genetic background. After four rounds of back-crossing, single heterozygous knockout virgins were crossed to single FM7 males to establish stocks. Unless indicated otherwise w^1118^ was used as the control strain throughout all experiments.

### Cell culture knockdowns and quantitative RT-PCR

dsRNAs were made using oligos OKH385/386 (anti CG34450, Supplemental Figure 2), OKH394/395 (anti CG34449, Figure 3A). S2 cells seeded in 6 well plates at 3×10^6^ cells/well/ml were treated with 12μg dsRNA in serum-free medium for 1h before 2ml of Schneider’s medium (GIBCO 21720) containing 10% FBS was added. After 5 days of knockdown cells were lysed and processed for SDS-PAGE and western blotting. Quantitative RT-PCR was performed using Maxima H Minus Reverse Transcriptase (Thermo Scientific EP0752), Maxima SYBG/ROX qPCR Master mix (Thermo Scientific K0223) and oligos OKH383/384 (PCR A, Supplemental Figure 2), OKH375/376 (PCR B, Supplemental Figure 2), OKH371/372 (PCR C, Supplemental Figure 2). Oligo sequences can be found in Supplemental Table 4.

### Stainings and antibodies

Immunostainings of cells or wing discs were performed as described previously [33]. Antibody against dZDHHC8 was raised in guinea pigs immunized with protein expressed in E. coli from pKH276 (used for immunostainings) or pKH278 (used for western blots). Other antibodies used were rat anti HA-tag (clone 3F10, Roche, 11 867 423 001), mouse anti actin (Hybridoma Bank, JLA20), mouse anti β-Galactosidase (Promega, Z378A), rabbit anti GFP (Torrey Pines Biolab TP401), rabbit anti scribble (kind gift from David Bilder), rabbit anti ERK1 (Cell Signaling 4695).

### Triglyceride measurements

Triglyceride levels were measured in L3 larvae as described previously [33].

### Clone size and wing measurements

GFP marked dZDHHC8 knockout clones and lacZ marked twin clones were generated by heat-shocking FRT18, GFP, KO/FRT18, lacZ female larvae for 45 min at 37°C three days prior wandering. Wing discs were stained with anti-GFP and anti-β-Galactosidase antibodies and imaged a Leica SP8 concofocal system with a 40x objective. Adult wings were imaged using an epifluorescent microscope with a 2.5x or 10x (for trichome density) objective. Clone size, wing compartment area as well as area of ectopic vein material were measured with ImageJ polygon selection tool.

### Acyl-biotinyl-exchange (ABE) assay

Expression of UAS-constructs was induced 24h prior to lysis by a 45min heat-shock at 37°C. The ABE assay was performed as described in [34]: Circa 160 L2 larvae were homogenised with a plastic pistil in 1.2ml lysis buffer containing 1.7% Triton X-100, 10mM N-ethylmaleimide (NEM), 2x proteinase inhibitor cocktail and 2mM PMSF and lysed for 15min on ice. After tissues were spun down, as an input sample 40μl lysate were mixed with 10μl 5x Lämmli and subsequently boiled for 5min at 95°C. Differing from the protocol published by Wan et al. [34] biotinylated proteins were finally pulled down using 15μl slurry of streptavidin coupled, magnetic dyna beads (Invitrogen 65001) and eluted in 30μl elution buffer. After elution samples were mixed with 15μl 5x Lämmli and boiled for 5min at 95°C.

### Staining of cellular compartment

The following constructs were expressed in S2 cells to mark cellular organelles: pMT-GFP-KDEL marking endoplamic reticulum, pMT-myc-fringe Golgi-tethered, pUAS-HA-Rab5 marking endosomes, pMT-dLamp1-GFP marking lysosomes.

## Acknowledgments

We thank the DKFZ Mass Spectrometry Core facility for mass spec. analyses, Marilena Lutz for technical help, and Natalia Becker from the DKFZ Biostatistics department for help with analyzing the mass spec data. Anti-actin antibody was obtained from the Developmental Studies Hybridoma Bank developed under the auspices of the NICHD and maintained by The University of Iowa.

---

**Supplemental Figure 1:**
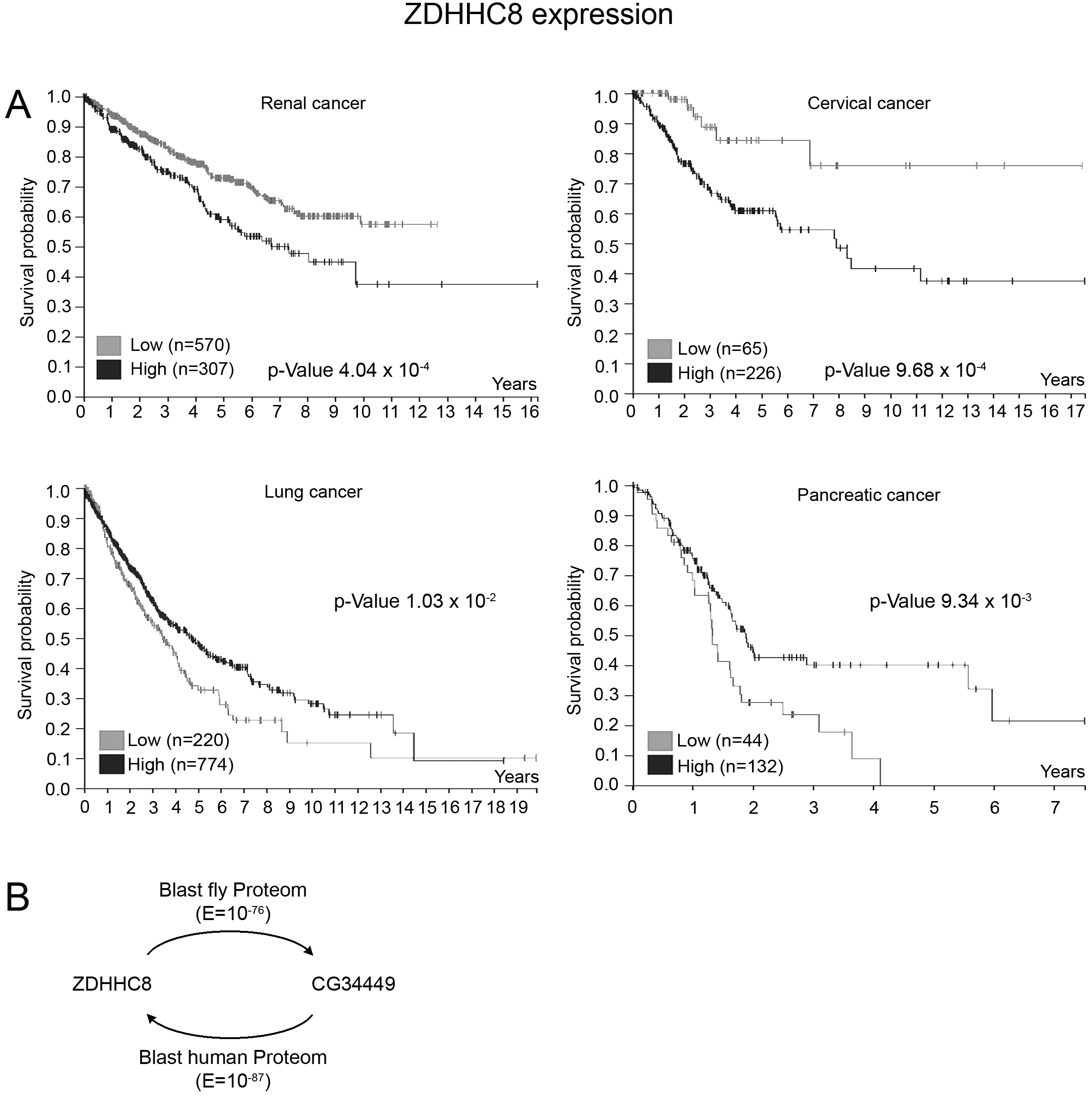
CG34449 is the Drosophila orthologue of human ZDHHC8. **(A)** ZDHHC8 expression levels correlate with increased or reduced cancer survival depending on cancer type. Kaplan-Meier plots (best separation, taken from The Human Protein Atlas using data from The Cancer Genome Atlas https://cancergenome.nih.gov) show a correlation between ZDHHC8 expression and patient survival for different cancer types. In renal (log-rank P value 4.04 x 10^−4^) and cervical cancer (log-rank P value 9.68 x 10^−4^) high ZDHHC8 expression correlates with decreased survival, whereas in lung (log-rank P value 1.03 x 10^−2^) and pancreatic cancer (log-rank P value 9.34 x 10^−3^) low ZDHHC8 expression correlates with decreased survival. **(B)** BLAST search using the Flybase BLAST server [35] of the Drosophila proteome using the protein sequence of human ZDHHC8 (NCBI Reference Sequence NP_037505.1) yields CG34449 as the top hit, with an E value of 10^−76^. Conversely, BLASTing the protein sequence of Drosophila CG34449 against the human proteome yields ZDHHC8 as the top hit with an E value of 10^−87^, identifying ZDHHC8 as the human orthologue of Drosophila CG34449.

**Supplemental Figure 2:**
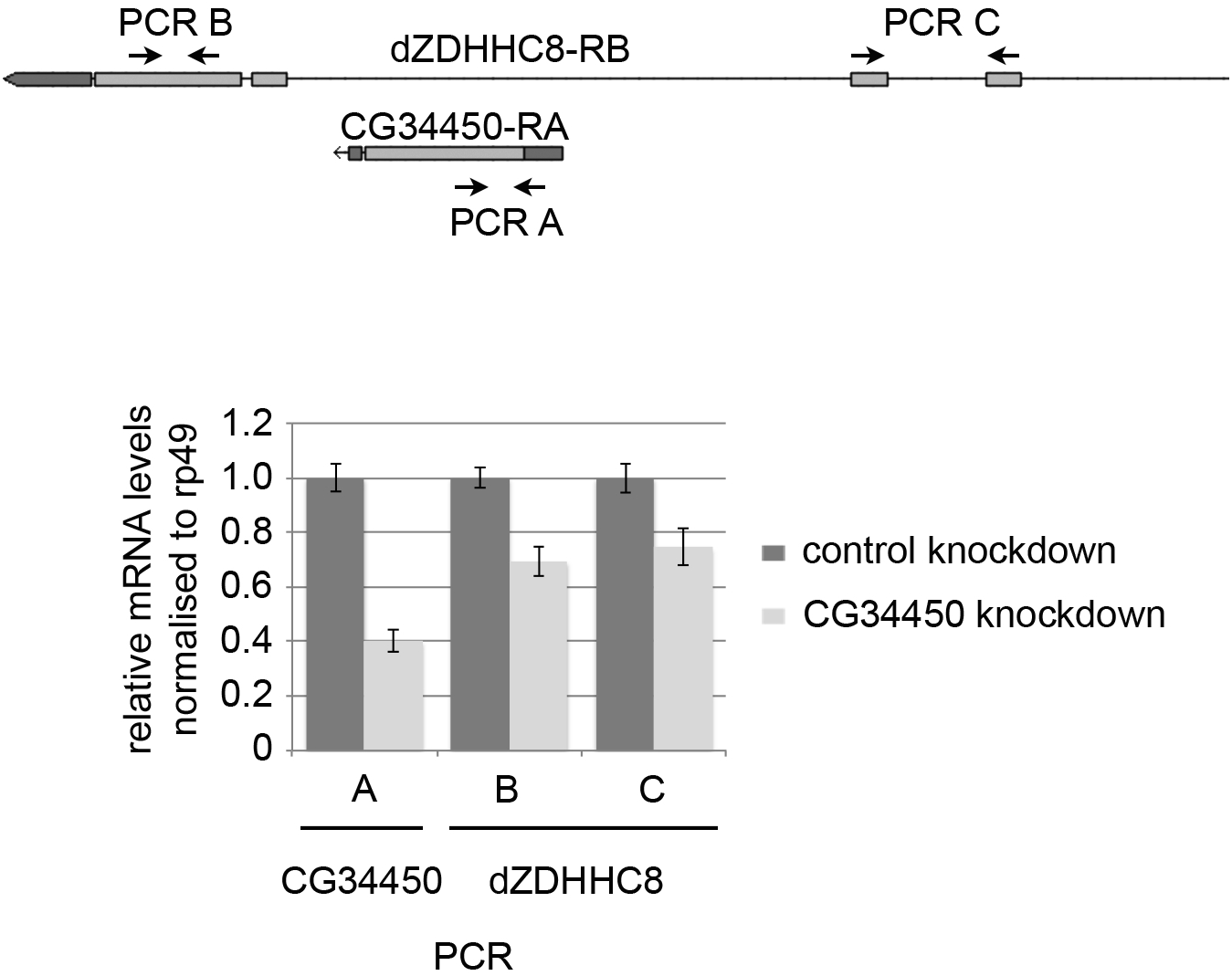
dZDHHC8 and CG34450 are transcriptionally linked. The expression of different dZDHHC8 exons (PCR B and C) was analysed by quantitative RT-PCR in control cells (dark grey bars) or cells with a CG34450 knockdown (light grey bars). When CG34450 is knocked down, transcript levels of dZDHHC8 also decrease suggesting dZDHHC8 and CG34450 are not separate genes.

**Supplemental Figure 3:**
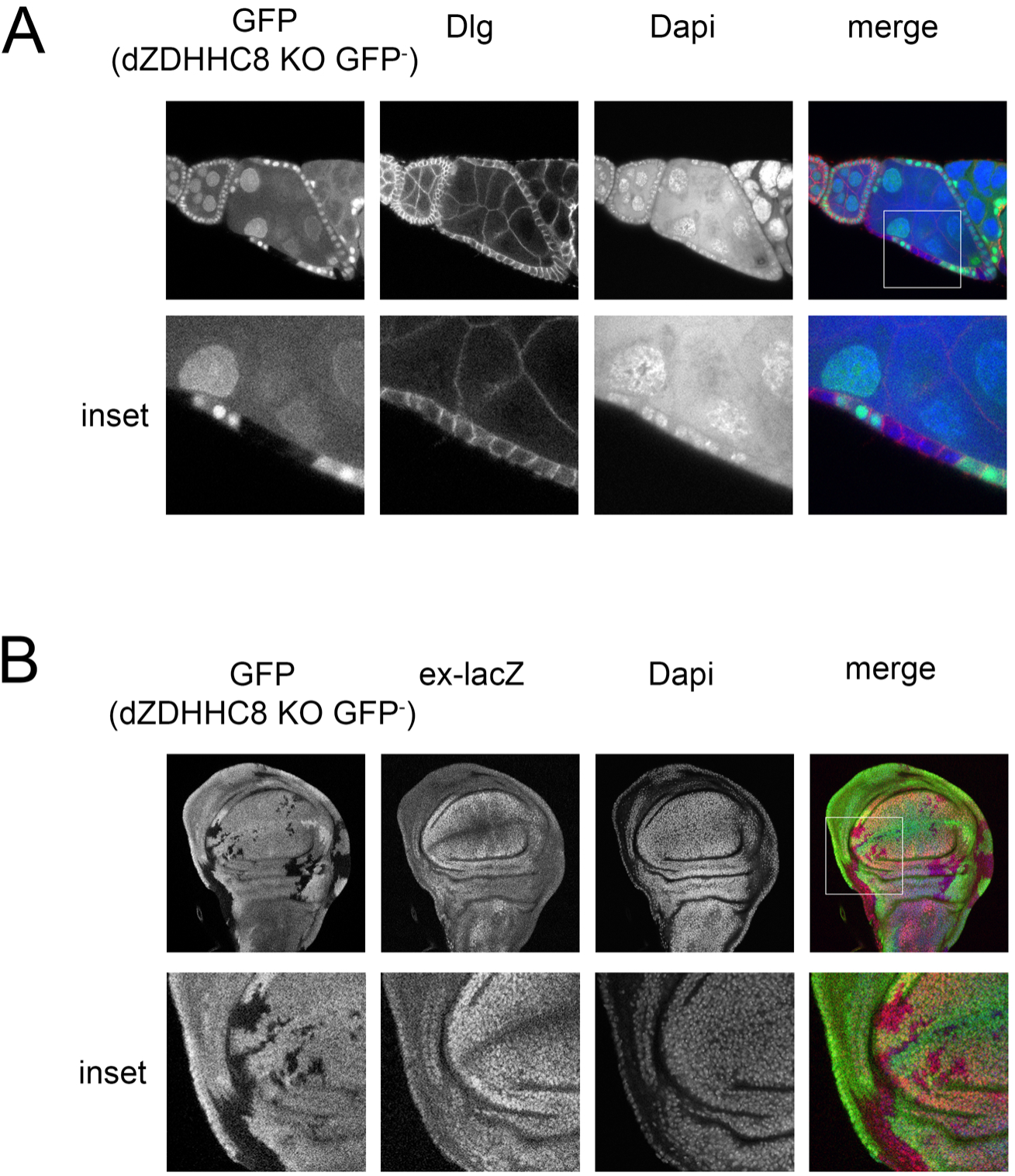
Scribble activity in dZDHHC8 mutant clones. **(A)** dZDHHC8 mutant follicle clones (GFP negative) were stained with an antibody detecting endogenous Dlg (red). Dlg localization is not affected in dZDHHC8 mutant clones. **(B)** Three day old dZDHHC8 knockout clones (GFP negative) in the wing disc were stained for the yorkie reporter ex-lacZ (red). Yorkie activity is not affected in dZDHHC8 mutant clones.

